# Visualization of breast cancer using contrast-enhanced optical coherence elastography based on tissue heterogeneity

**DOI:** 10.1101/2024.10.09.617341

**Authors:** Jiayue Li, Ken Y. Foo, Rowan W. Sanderson, Renate Zilkens, Mireille Hardie, Laura Gale, Yen L. Yeow, Celia Green, Farah Abdul-Aziz, Juliana Hamzah, James Stephenson, Ammar Tayaran, Jose Cid Fernandez, Lee Jackson, Synn Lynn Chin, Saud Hamza, Anmol Rijhumal, Christobel M. Saunders, Brendan F. Kennedy

## Abstract

By mapping the mechanical properties of tissue, elastography can improve identification of breast cancer. On the macro-scale, ultrasound elastography and magnetic resonance elastography have emerged as effective clinical methods for the diagnosis of tumors. On the micro-scale, optical coherence elastography (OCE) shows promise for intraoperative tumor margin assessment during breast-conserving surgery. Whilst several OCE studies have demonstrated strong potential, the mechanical models used require the assumption of uniaxial stress throughout the sample. However, breast tissue is heterogeneous and contains compressible features (*e*.*g*., ducts and blood vessels) and collagen-rich fibrotic features (*e*.*g*., stroma). This heterogeneity can invalidate the assumption of uniaxial stress and reduce the accuracy of OCE, often making it challenging to interpret images. Here, we demonstrate a new variant of OCE based on mapping the Euler angle, *i*.*e*., the angle between the principal compression and the loading axis induced by tissue heterogeneity, which removes the assumption of uniaxial deformation. This is enabled by a hybrid three-dimensional (3-D) displacement estimation method that combines phase-sensitive detection and complex cross-correlation, providing access to the 3-D displacement and 3-D strain tensor on the micro-scale. We demonstrate this new OCE technique through experiments on phantoms and 10 fresh human breast specimens. Through close correspondence with histology, our results show that mapping the Euler angle provides additional contrast to both optical coherence tomography and a current OCE technique in identifying cancer. Mapping the Euler angle in breast tissue may provide a new biomarker for intraoperative tumor margin assessment.

## I. Introduction

BREAST cancer is either the first or second leading cause of female cancer death in 95% of countries [1]. Breast-conserving surgery is the most common surgical treatment for early-stage breast cancer. In this procedure, the aim is to remove malignant tissue while achieving a good cosmetic outcome for patients [2]. However, in 20–30% of cases, patients undergo re-excision due to inadequate margins identified in post-operative histology [3], [4]. Re-excision increases risk of infection, treatment costs, and psychological burden to patients. Accurate intraoperative assessment of tumor margins would allow surgeons to remove the tumor more completely during the initial surgery, potentially reducing the re-excision rate. A number of intraoperative diagnostic techniques have been developed to aid in tumor margin assessment [5], including histopathological techniques, such as frozen section analysis and imprint cytology, and medical imaging techniques, such as specimen radiography, ultrasonography, and magnetic resonance imaging. However, none of these methods have been widely adopted in clinical settings, as they are limited by the tradeoffs between speed, cost, spatial resolution, and diagnostic accuracy [5], [6]. More recently, optical imaging techniques providing micro-scale resolution, such as Raman spectroscopy and optical coherence tomography (OCT), have shown promise in intraoperative tumor margin assessment. However, Raman spectroscopy has long acquisition times, typically 12–24 minutes per margin [7], and while OCT can be performed in intraoperative timeframes, it often exhibits low contrast between cancerous and stromal tissues [8].

Beyond optical contrast, the modification of tissue during tumor initiation and progression provides mechanical contrast between malignant and benign tissues [9]–[12]. Imaging tissue mechanical properties on the micro-scale has the potential to improve intraoperative identification of breast cancer. Optical coherence elastography (OCE), particularly compression OCE, has shown promise as a diagnostic tool for intraoperative tumor margin assessment [13]–[16]. In this technique, tissue deformation induced by a compressive load is tracked using OCT with sub-nanometer displacement sensitivity [17]. A mechanical model is then used to convert the measured deformation into micro-scale elasticity (quantified by Young’s modulus), which is mapped into three-dimensional (3-D) elastograms [18].

In OCE, as is the case in all elastography techniques, assumptions are made on the nature of tissue deformation to simplify the mechanical models used to estimate elasticity. In compression OCE, the sample is assumed to be linear elastic and isotropic [19], [20]. Furthermore, stress is assumed to be uniaxial and uniform in depth. In this case, by measuring compressive stress and compressive strain, which are both negative by convention, local elasticity can be calculated as the ratio of the normal axial component of the stress tensor (*i*.*e*., axial stress) at the sample surface to the normal axial component of the strain tensor (*i*.*e*., axial strain) within the sample. The surface stress may be measured using, for example, a load cell [21], [22] or a pre-characterized compliant layer [23], [24]. Whilst elasticity imaging often provides high contrast in differentiating tumor from surrounding tissues, tissue heterogeneity can invalidate the assumption of uniaxial compressive stress and strain throughout the sample [25]. For instance, in breast cancer, nests of tumor cells and intermixed immature stroma result in mechanical heterogeneity that can cause the principal planes, where there is highest compression, to rotate and deviate from the axis of uniaxial loading, resulting in adjacent regions of negative (compressive) and positive (tensile) axial strain [15], [26]. In addition, tissue features, such as ducts [25], vessels [27], and fibrotic stroma [13], [25] exhibit structural heterogeneity, which can also cause positive axial strain. Importantly, under the assumption of uniaxial stress in compression OCE, the measured negative surface stress and positive strain in some regions of the sample are converted to negative elasticity, which is not physically meaningful [13], [16], [25]. These effects highlight the limitations of oversimplified models of complex tissue.

Mapping the deviation from uniaxial compression in the local strain tensor could offer a new contrast mechanism to identify regions of tumor. To assess this potential, we implemented a hybrid 3-D displacement method that employs complex cross-correlation [28] combined with phase-sensitive detection [17]. This provides access to the full 3-D strain tensor, enabling the angle between the principal compression and the loading axis (*z*-axis), *i*.*e*. the Euler angle of principal compression, to be determined. In this study, we demonstrate Euler angle imaging through experiments on phantoms and 10 freshly excised human breast specimens. We compare Euler angle imaging with co-registered histology, OCT, and compression OCE, *i*.*e*., elasticity imaging. We show that Euler angle imaging provides complementary and, in many cases, superior contrast to both OCT and elasticity in breast tumor and surrounding stroma. This enables a more comprehensive characterization of human breast tissue and may facilitate improved intraoperative tumor margin assessment.

## II. Background

### A. Mechanical model of compression OCE

In compression OCE, to minimize the effects of viscoelasticity, adequate time (>100 seconds [29]) is given after a pre-strain is applied to the sample to yield a steady state. A micro-scale load (actuation) is then introduced to deform the sample. The applied load and the resulting deformation can be described in terms of stress and strain tensors, which enable elasticity to be computed from the constitutive equation that links the two tensors [30]. If the sample exhibits linear elastic behavior, which is valid for soft tissues under low applied strain [31], [32], and if the deformation is infinitesimally small relative to the spatial coordinates of the body, the stress tensor, also known as the Cauchy stress tensor, *σ*, is given by [33]

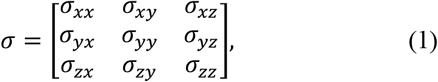

and the infinitesimal Cauchy strain tensor, *ε*, is given by [33]

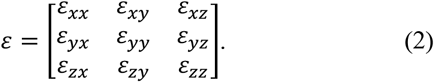

For each component of both tensors, the first subscript denotes the normal plane on which the component is exerted or induced, and the second denotes the direction of the component. The constitutive equation, also known as the generalized Hooke’s law, is given by [30]

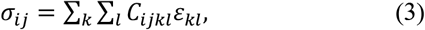

where *C*_*ijkl*_ represents a 4^th^ order elasticity tensor, which can be written as a 2^nd^ order tensor (in Voigt notation) with 36 elastic constants. Under the further assumption that the sample is mechanically isotropic, (3) can be simplified to

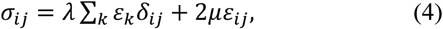

where the elastic constants are reduced to the two Lamé constants, *λ* and *μ* (where *μ* is also known as shear modulus), *δ*_*ij*_ is the Kronecker delta function (*δ*_*ij*_ = 1 if *i* = *j*, and 0 otherwise), and ∑_*k*_ *ε*_*k*_ represents the summation of normal strain components. Importantly, *λ* and *μ* can be defined in terms of Young’s modulus, *E*, and Poisson’s ratio, *υ*, two commonly used mechanical properties in biomechanics, allowing (4) to be rewritten as

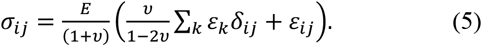

Assuming that the sample is deformed uniaxially under the applied uniaxial stress in *z, i*.*e*., *i* = *j* = *z, σ*_*ij*≠*zz*_ = 0, and ∑_*k*_ *ε*_*k*_ = *ε*_*zz*_(1 − 2υ) for an isotropic medium, (5) can be rearranged and written as

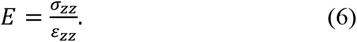

In compression OCE, the axial component of displacement, *i*.*e*., *u*_*z*_, is generally measured using phase-sensitive OCT with sub-nanometer displacement sensitivity [17] and *ε*_*zz*_ is then calculated as the gradient of *u*_*z*_ with depth [26]. By placing a compliant layer with known stress-strain response on the sample surface, the measured axial strain in the layer is related to axial stress at the sample surface [23]. Assuming that surface stress is uniaxial and uniform in depth, elasticity at each spatial location can be estimated as the ratio of local axial surface stress and local axial sample strain.

### B. Strain tensor transformation

As depicted in Fig. 1, the components of the strain tensor in (2) can be transformed to a new coordinate system, in which all shear strain components are zero by solving the eigenvalue problem [34]

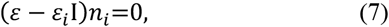

*where* I *is the identity matrix, ε*_*i*_ and *n*_*i*_ are eigenvalues and eigenvectors, respectively, corresponding to principal strains, *ε*_1_, *ε*_2_, and *ε*_3_ (*ε*_1_ < *ε*_2_ < *ε*_3_), and principal axes, *n*_1_, *n*_2_, and *n*_3_. The minimum (*ε*_1_) and maximum (*ε*_3_) principal strains and the corresponding axes (*n*_1_ and *n*_3_) represent the amplitude and direction of the greatest compression and tension, respectively. The angle differences between the principal axes and the initial axes are defined as the Euler angle of principal strains [35], *α, β*, and *γ*. Specifically, *α* is defined as the Euler angle of principal compression, which is the parameter most relevant to compression OCE and is referred to simply as Euler angle from here on.

**Fig. 1.**
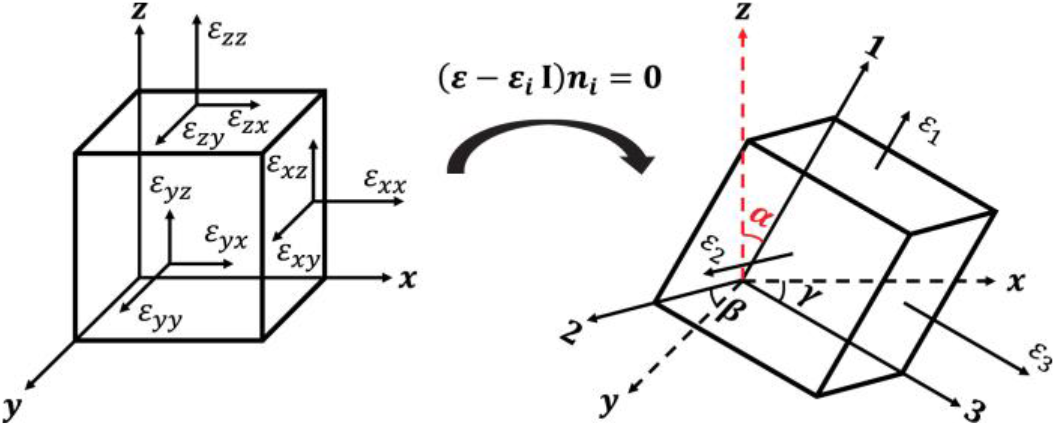
Transformation of the strain tensor, *ε*, to the coordinate of principal strain, *ε*_*i*_.The Euler angle is denoted as *α*.

## III. Materials and Methods

### A. Phantom fabrication

To illustrate Euler angle mapping, we first fabricated a uniform phantom, phantoms containing stiff inclusions with varying elasticity, and a phantom comprising a cylindrical cavity. The phantoms were fabricated using two-component vulcanizing silicone elastomers, namely Silpuran (Wacker Chemie AG, Germany) and RT601 (Wacker Chemie AG, Germany), mixed with PDMS oil (AK50, Wacker Chemie AG, Germany) to vary the elasticity of the phantoms. The phantoms were cylindrical with thickness of 3 mm and diameter of 6 mm, respectively.

The uniform phantom and the soft bulk of the inclusion phantoms were made from the same material with Young’s modulus of 47.6 kPa, characterized using the same protocol for uniaxial compression testing as described in a previous study [36]. The three stiff inclusions are cubes with a side length of 1 mm, embedded ∼500 µm below the surface of the soft bulk, with Young’s moduli of 67.4 kPa, 147.4 kPa, and 236.1 kPa, respectively, resulting in a mechanical contrast ratio in the inclusion phantoms of 1.4, 3.1, and 5.0, respectively.

The cavity phantom is a uniform phantom containing a compressible, hollow, cylindrical channel located close to the central axis. To fabricate this phantom, a mold with holes of diameter ∼600 µm located on the opposite sides of the walls was printed using a stereolithography 3-D printer (Form 2, Formlabs, USA). A 23-gauge needle was inserted through the holes before casting the silicone mixture into the 3-D printed mold. The same silicone was used as for the uniform phantom. After curing, the needle was removed, resulting in a 6 mm long cylindrical channel, ∼500 µm below the surface. To introduce optical scattering to the phantoms, titanium dioxide (TiO_2_) particles (refractive index = 2.3, Sigma-Aldrich, Germany) were added and evenly mixed in the silicone at a concentration of 2 mg/ml prior to curing. In the inclusion phantoms, TiO_2_ particles were added to the bulk and inclusion in concentrations of 2 mg/ml and 5 mg/ml, respectively.

### B. Tissue preparation, histology, and co-registration

In this study, 10 patients undergoing mastectomy were recruited from two sites in Western Australia, Fiona Stanley Hospital and Sir Charles Gairdner Hospital, with ethics approved by South Metropolitan Health Service Human Research Ethics Committee (PRN: RGS0000003726) and Sir Charles Gairdner and Osborne Park Health Care Group Human Research Ethics Committee (PRN: RGS0000001694), respectively. Ten specimens were obtained with informed consent and were imaged within an hour of excision.

After data acquisition, each specimen was inked for orientation, placed in a cassette, and fixed in 10% neutral-buffered formalin for at least one day before histology processing. Paraffin embedding was performed prior to sectioning and staining with hematoxylin and eosin (H&E), following the standard histology protocols used at the hospital. The H&E histology slides were then digitized using a slide scanner (Axioscan 7, Zeiss, Germany) and co-registered with *en face* OCT images and elastograms following protocols described previously [37]. Each histology slide was reviewed and annotated by a pathologist to identify tissue components.

### C. Experimental setup

Compression OCE measurements were performed using a fiber-based spectral-domain OCT system (Telesto 320; Thorlabs Inc., USA) with a central wavelength of 1300 nm and a 3-dB bandwidth of 170 nm. The measured axial and lateral resolution (full width at half maximum (FWHM)) is 4.8 µm (in air) and 7.2 µm, respectively, for the objective lens used (LSM03; Thorlabs Inc., USA). A 4 mm thick glass window (Edmund Optics Inc., USA) with a diameter of 75 mm was rigidly affixed to a ring actuator (Piezomechanik GmbH, Germany) with an internal aperture of 65 mm and a maximum stroke of 10 µm, enabling imaging and micro-scale compression to be introduced from the same side of the sample [25].

A ∼500 µm thick translucent silicone layer (Elastosil P7676, Wacker Chemie AG, Germany) with Young’s modulus of 20.2 kPa was placed between the window and the sample to map stress at the sample surface [23]. The OCT system was configured in common-path mode where the layer-window interface provided the reference reflection [38]. A motorized vertical translation stage (MLJ150; Thorlabs Inc., USA) was used to bring the sample and layer into contact with the window. A pre-strain (*i*.*e*., a bulk strain prior to actuation) of ∼5–20% was then applied to ensure uniform contact between the window, layer, and tissue sample. At the given pre-strain, the actuator was driven collinearly with the OCT beam and synchronized with the OCT B-scan pairs, such that alternate B-scans were acquired in different loading states. Each B-scan comprises 2,000 A-scans in *x*. For mechanically isotropic phantoms, 3-D scans comprising 200 B-scan pairs (unloaded and loaded) were acquired in *y*. The uniform phantom was scanned over a 7 mm × 0.7 mm (*xy*) field-of-view (FOV). The inclusion phantoms and the cavity phantom were scanned over a FOV of 4 mm × 0.4 mm (*xy*). Each dataset was acquired in 11.2 seconds. For the results acquired on phantoms, the middle 100 B-scan pairs were spatially averaged to improve strain sensitivity. 3-D scans on freshly excised breast tissue comprising 2,000 B-scans pairs were acquired over a FOV of 10 mm × 10 mm (*xy*) at an acquisition time of 112 seconds. For the phantom scans, the contact boundaries were lubricated with PDMS oil to reduce surface friction. For human breast tissue, saline was applied at the layer-tissue boundary both to reduce surface friction and to keep the tissue hydrated during scanning.

### D. Signal processing in compression OCE

As described in Section II.B, the Euler angle is derived from the 3-D strain tensor, which can be calculated as the spatial derivative of the 3-D displacement,

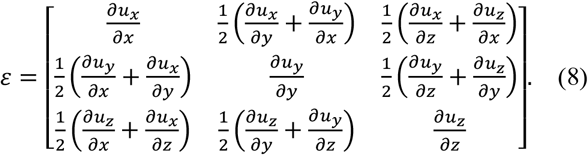

To estimate tissue displacement and the corresponding strain tensor components, a hybrid 3-D displacement method was developed that combines phase-sensitive detection [17], for the estimation of *u*_*z*_, with a non-iterative, complex cross-correlation method [28], for the estimation of lateral displacement components, *i*.*e*., *u*_*x*_ and *u*_*y*_. In phase-sensitive detection, *u*_*z*_ was estimated from the phase difference between an unloaded-loaded B-scan pair acquired at the same spatial location [38]. The lateral and transverse displacements were estimated from the set of equations defined by the amplitude of complex correlation coefficients (ACCC) using the complex cross-correlation method with nanometer-scale displacement sensitivity [27]. A detailed description can be found elsewhere [27], [39]. Briefly, the ACCC was calculated between the unloaded complex OCT volume and a set of the loaded complex OCT volumes including the original and bi-directional copies, shifted by one voxel in *x, y*, and *z*, respectively, over a local neighborhood window defined by a Gaussian convolution kernel. As the OCT speckle size can be approximated as √2× the OCT resolution [40], [41], we chose a displacement kernel that is 3× the OCT resolution, *i*.*e*., ∼22 μm × 22 μm × 14 μm (*xyz*) to ensure high displacement resolution, whilst still including a sufficient number of speckles to maintain high displacement accuracy and sensitivity [27], [42]. To reduce the effect of speckle and optical noise, we smoothed displacement volumes using a 3-D Gaussian kernel with a size of 72 μm × 72 μm × 48 μm (*xyz*).

For strain estimation, weighted least squares (WLS) linear regression was used to estimate each component of the strain tensor over a fitting range of 120 μm (FWHM) [43]. Then, the spatial derivatives of axial displacement, *i*.*e*., 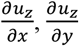, and 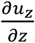, were estimated from the phase difference-derived axial displacement, and the remaining components, 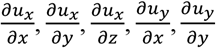, and 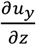, were estimated from ACCC-derived lateral and transverse displacement. The isotropic strain resolution (FWHM) was ∼140 μm (*xyz*), calculated as the convolution of the OCT system resolution and the FWHM of the signal processing used, *i*.*e*., 3-D Gaussian smoothing and 1-D WLS linear regression [44]. From the full strain tensor, the principal compressive strain and the Euler angle of principal compression were then determined from solutions to the “eigenvalue problem” in (7), as described in Section II.B. Sample elasticity is calculated as the ratio of local axial stress at the sample surface to local axial strain within the sample, by assuming the surface stress is uniaxial and is distributed uniformly in depth [23]. The resultant elasticity resolution (FWHM) was ∼7.2 μm × 7.2 μm × 140 μm (*xyz*).

## IV. Results

### A. Phantoms

Results obtained from the phantoms are shown in Fig. 2. Figure 2a presents results of the uniform phantom, showing the cross-sectional (*xz*) OCT B-scan, overlaid with the axial and lateral (*x*-) components of the displacement field, along with the corresponding axial strain, elasticity, and Euler angle. For all elastography images in Fig. 2, the region of the homogeneous layer was masked. With the system configured in common-path mode, under compression, the axial displacement is small close to the layer-window interface, and increases in depth, indicating negative axial strain, *i*.*e*., compressive strain. For nearly incompressible materials to conserve their volume, axial compression is accompanied by lateral expansion. This is illustrated in Fig. 2a by the lateral displacement increasing gradually for locations away from the central axis. As expected, the corresponding elastograms in Fig. 2a demonstrate uniform distribution of both strain and elasticity and, furthermore, Euler angle is close to zero.

**Fig. 2.**
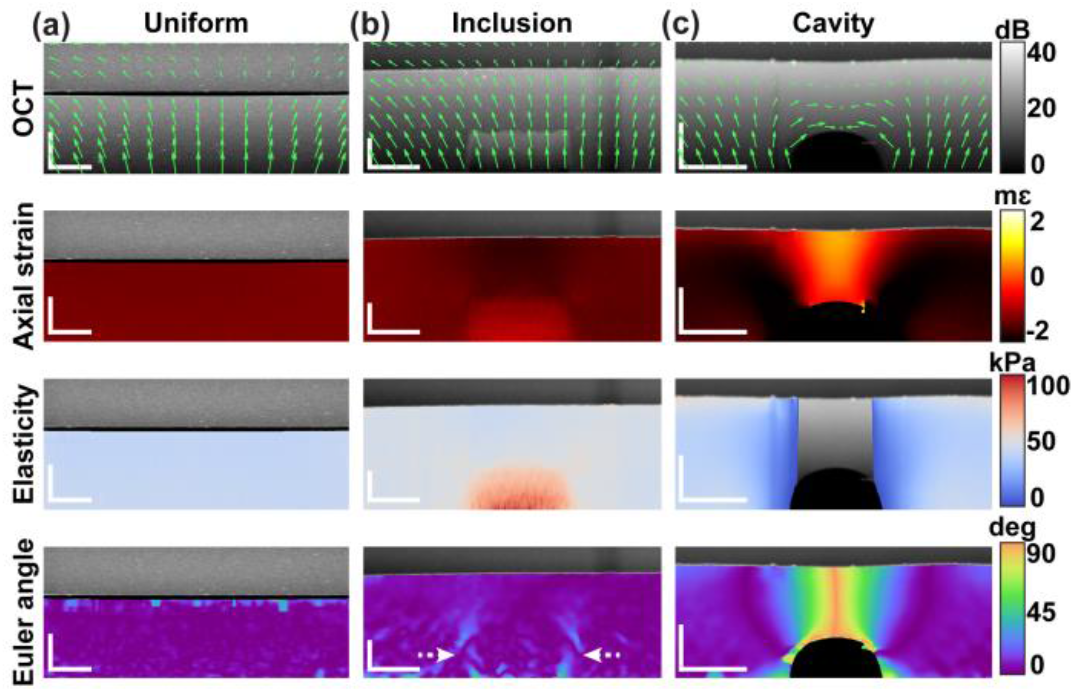
Phantoms results. Cross-sectional OCT B-scans (*xz*) overlaid with displacement vectors and the corresponding elastograms of strain, elasticity, and Euler angle of principal compression of (a) a uniform phantom, (b) an inclusion phantom, and (c) a cavity phantom, respectively. Elastograms are overlaid on OCT images. Scale bars represent 500 µm.

The inclusion phantom is presented in Fig. 2b. As in the uniform phantom, increased lateral displacement in regions away from the central axis indicates the incompressibility of both the stiff inclusion and the soft bulk. The stiff inclusion is clearly visible in both strain and elasticity images. There is a noticeably lower strain (*i*.*e*., higher compression) in the soft bulk observed in the region above the stiff inclusion than in the region adjacent to the inclusion. This is expected as the region above the inclusion is closer to the top rigid window and, thus, is axially confined between the window and inclusion. This creates a non-uniform stress distribution, where a higher magnitude of stress is present in the region above the inclusion than in the region adjacent to the inclusion. The non-uniform stress is accounted for in the elasticity estimation by mapping surface stress at the layer-sample interface [23], as illustrated by the relatively uniform elasticity in the bulk in the corresponding elasticity image. In the Euler angle image, the inclusion boundaries are delineated with higher Euler angle at the boundaries (indicated by white dashed arrows) than in the regions away from the boundaries. This is attributed to mechanical coupling between the inclusion and the surrounding bulk material, leading to non-uniaxial and non-uniform stress distribution at the inclusion boundaries, particularly in the bulk close to the inclusion vertices with localized stress that has the greatest magnitude caused by the abrupt change in geometry. This stress variation induces increased shear strain at the boundaries, which, in turn, causes the direction of principal compressive strain to deviate from the applied loading axis.

As observed in Fig. 2c, the cavity phantom exhibits a distinct deformation pattern, where the overlaid displacement vectors change direction sharply in the region above the cavity, converging towards the cavity, rather than diverging from the central axis as in the case of the uniform phantom (Fig. 2a) and the inclusion phantom (Fig. 2b). This indicates that lateral compression (contraction) and axial tension (expansion) occur in the region above the cavity due to the nearly incompressible nature of the material under compression. This suggests that the hollow cavity beneath reduces in volume under compression [27], which is supported by the corresponding axial strain elastogram depicting positive axial strain in the region above the cavity. Furthermore, the positive axial strain within the sample, and negative axial stress estimated from strain in the compliant layer results in negative elasticity, which invalidates the mechanical model of compression OCE. This negative elasticity is masked in Fig. 2c. As highlighted in the corresponding Euler angle image, the Euler angle approaches 90º above the cavity, suggesting reduction in volume of the underlying compressible feature, which causes rotation of the principal compression axis orthogonal to the *z*-axis (loading).

### B. Inclusion phantoms with increased mechanical contrast

Figures 3a and 3b show OCT and elasticity B-scans (*xz*) of inclusion phantoms with mechanical contrast ratios (CR = *E*_inclusion_ : *E*_matrix_) of 1.4, 3.1, and 5.0, respectively. The plots of OCT intensity and elasticity as a function of *x*, ∼255 µm below the inclusions (indicated by the black dashed lines), are presented in Fig. 3d and 3e, respectively. The plots were generated by averaging measurements over ∼70 µm in depth. As expected, the elasticity contrast increases as the inclusion elasticity increases, while the OCT contrast remains relatively constant. For validation, in the elasticity plots (Fig. 3e), dashed lines indicate the elasticity of the silicone materials characterized using uniaxial compression testing. The plots show that the accuracy of quantitative OCE measurements in both the bulk and inclusion is reduced as the mechanical contrast increases. This is likely due to an increase in the stress-shielding effect [45], *i*.*e*., stress non-uniformity, in the bulk regions adjacent to the sides of the inclusions with greater Young’s modulus. In addition, Fig. 3c shows that the contrast in the Euler angle images increases as the mechanical contrast increases. This is likely due to increased mechanical coupling between the inclusion and matrix materials under compression, and increased maximum stress magnitude localized in the vicinity of the inclusion vertices. This is highlighted in the plots of Euler angle, ∼35 µm above the inclusions, in Fig. 3f, demonstrating the ability to delineate mechanical heterogeneity by mapping Euler angle.

**Fig. 3.**
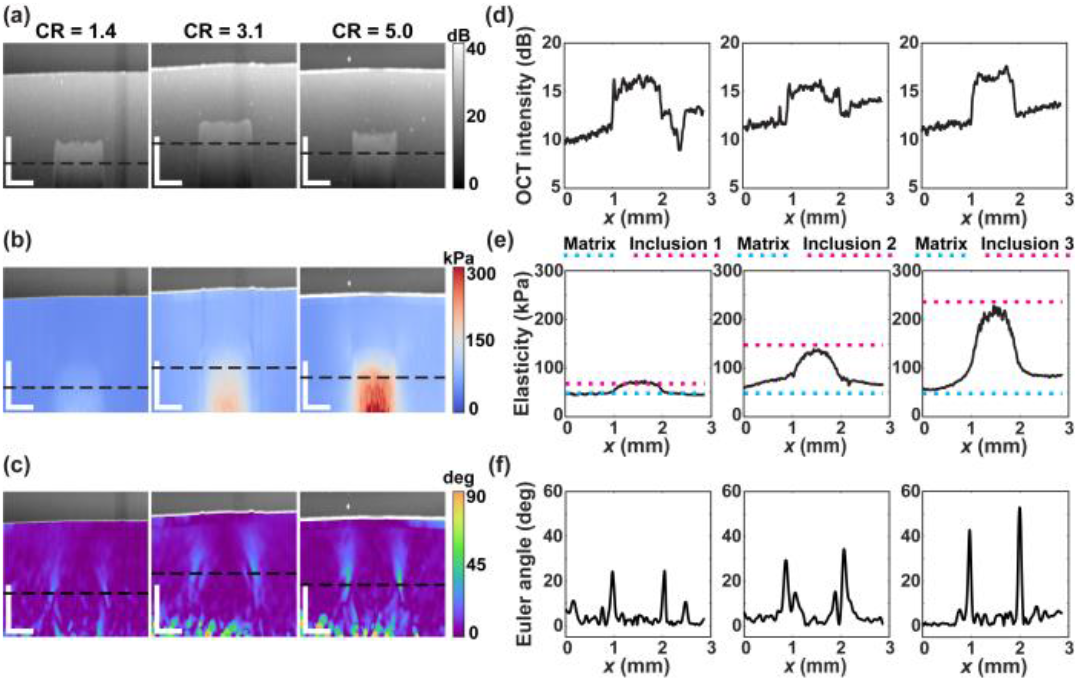
Inclusion phantoms with increasing mechanical contrast. (a) OCT B-scans (*xz*) and corresponding elastograms of (b) elasticity and (c) Euler angle of inclusion phantoms with mechanical contrast ratio (CR) of 1.4, 3.1, and 5.0, respectively. 1-D plots are shown for (d) OCT intensity and (e) elasticity as a function of *x*, ∼255 µm below the inclusions, corresponding to black dashed lines in (a) and (b). The blue and magenta dashed lines in (e) represent the bulk elasticity of the matrix and inclusion materials characterized by uniaxial compression testing. (f) The 1-D plots of Euler angle as a function of *x*, ∼35 µm above the inclusion, corresponding to black dashed lines in (c). Scale bars represent 500 µm.

### C. Visualization of human breast tissue using OCT, elasticity, and Euler angle of principal compression

In Figs. 4–6, to demonstrate the potential of Euler angle imaging for visualization of breast cancer, we present example Euler angle images of freshly excised human breast tissue, along with the co-registered histology, and corresponding OCT and elasticity.

**Fig. 4.**
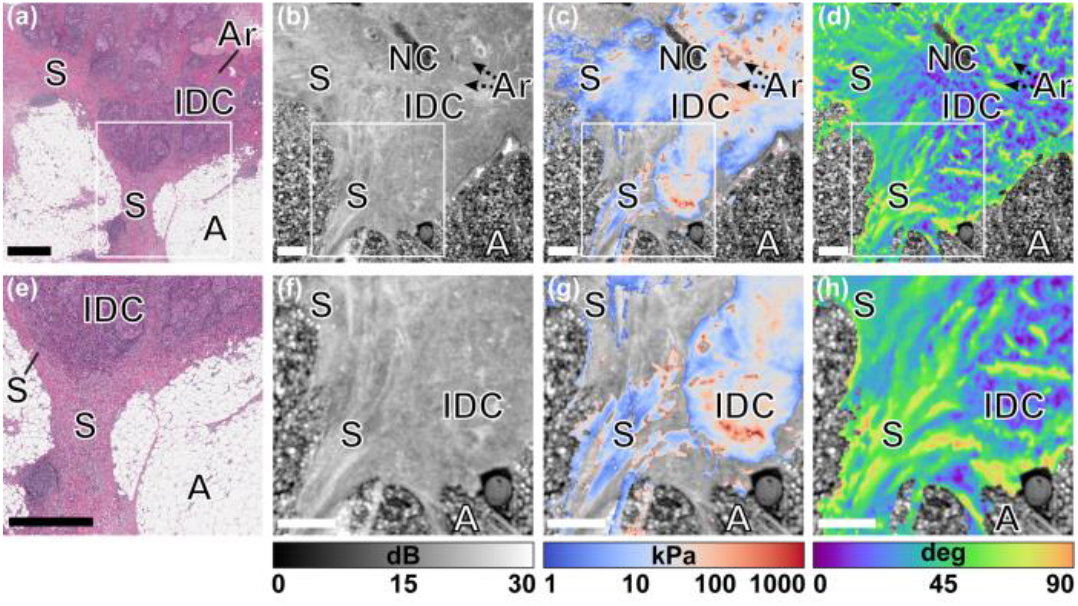
A mastectomy specimen containing Grade 2 IDC. (a) Histology image of the specimen. The co-registered *en face* (*xy*) images of (b) OCT, (c) elasticity, and (d) Euler angle, ∼200 µm below the tissue surface. The white boxes shown in (a)–(d) indicate the magnified regions presented in (e)–(h). S: stroma; NC: non-contact area; Ar: artery; IDC: invasive ductal carcinoma; A: adipose tissue. Scale bars represent 1 mm.

Figure 4 shows results from a mastectomy specimen containing Grade 2 invasive ductal carcinoma (IDC). The histology, *en face* OCT, elasticity, and Euler angle images are presented in Figs. 4a–4d, respectively. The binary segmentation algorithm [37] used to generate images with elastograms overlaid on OCT images identifies regions of low OCT intensity, mainly in adipose tissue, and masks these pixels from the elastograms. The OCT image in Fig. 4b depicts a heterogeneous pattern of IDC and highly scattering thin strands of fibrotic stroma to the left of IDC and adjacent to adipose tissue. This is further highlighted in Fig. 4f, which shows a magnified region marked by a white box in Fig. 4b. The corresponding elasticity in Fig. 4g shows high elasticity in IDC, which is likely due to densely packed tumor cells that invoke a desmoplastic response at the invasive front of the tumor [46]. By contrast, adjacent stroma exhibits low and relatively uniform elasticity. In addition, compressible features, such as the artery in this specimen, can reduce in volume, resulting in positive axial strain in the region surrounding the tubular structures, which leads to negative elasticity, as indicated by the black dashed arrows in Fig. 4c and Fig. 4d. The Euler angle image (Fig. 4d) shows that the localized Euler angle approaches 90º in regions where the elasticity is negative. Furthermore, the Euler angle accentuates the boundaries between stroma and IDC, showing that the Euler angle tends to be high in stroma and low in IDC, which suggests that, in this example, IDC exhibits strain more closely aligned to the normal (*z*) axis. Improved contrast can also be observed in the magnified region of the Euler angle image in Fig. 4h. Correspondence with the OCT image (Fig. 4f) suggests that the contrast provided by the Euler angle enables the orientation of the stromal bands adjacent to IDC to be delineated. We note that the unique pattern of localized Euler angle in stromal bands is caused by high shear strain measured in experiment, suggesting relative sliding between adjacent connective tissue under compression. In the corresponding elasticity image (Fig. 4g), large regions present negative elasticity and are therefore masked from the image, due to the existence of local positive axial strain in the stromal bands.

Figure 5 presents the results of a mastectomy specimen containing Grade 3 invasive pleomorphic lobular carcinoma (IPLC). The histology image in Fig. 5a reveals invasive cancer surrounded by stroma with clusters of adipose tissue. The co-registered *en face* OCT, elasticity, and Euler angle images are presented in Figs. 5b–5d, respectively. The OCT image in Fig. 5b, and the corresponding magnified image in Fig. 5f, show that solid aggregates of tumor cells, appearing as regions of low OCT intensity, can be differentiated from the surrounding stroma with relatively higher OCT intensity, and adipose tissue appearing as small, discrete regions of low intensity. The corresponding elasticity in Fig. 5c shows that IPLC exhibits higher elasticity than the surrounding stroma, despite small portions of invalid elasticity around the boundaries of the tumors being masked from the image, which correspond to elevated Euler angle in Fig. 5d. This is highlighted in a magnified region containing tumor and stroma in Figs. 5g and 5h, respectively, corresponding to the region marked by white boxes in Figs. 5c and 5d. Furthermore, through close correspondence with histology in Fig. 5e, it is clear that the Euler angle (Fig. 5h) delineates the boundary of the tumor with lower Euler angle than the adjacent stroma, providing additional contrast to OCT (Fig. 5f) and elasticity (Fig. 5g).

**Fig. 5.**
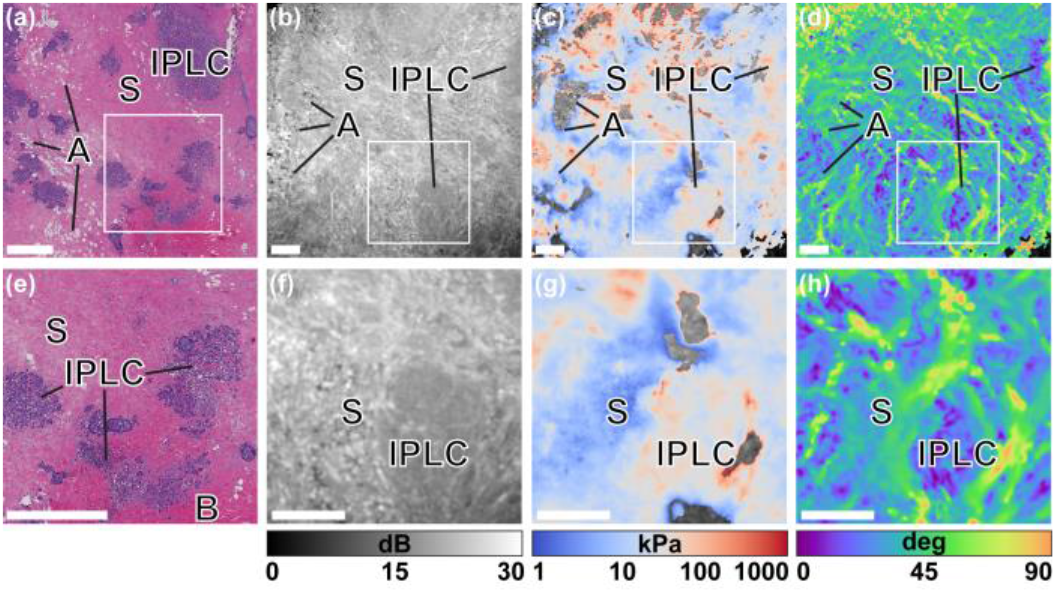
A mastectomy specimen containing Grade 3 IPLC. (a) Histology image of the specimen. The co-registered *en face* (*xy*) images of (b) OCT, (c) elasticity, and (d) Euler angle, ∼140 µm below the tissue surface. The white boxes shown in (a)–(d) indicate the magnified regions shown in (e)–(h), respectively. S: stroma; IPLC: invasive pleomorphic lobular carcinoma; A: adipose tissue, B: blood cells due to hemorrhage. Scale bars represent 1 mm.

Figure 6 presents the results of a mastectomy specimen containing Grade 2 IDC and intermediate grade ductal carcinoma *in situ* (DCIS). The histology image in Fig. 6a shows increased tumor proliferation leading to high density of tumor cells intermixed with stromal tissue. The co-registered *en face* OCT, elasticity, and Euler angle images are shown in Figs. 6b– 6d, respectively. Compared to the OCT image (Fig. 6b), which exhibits relatively low contrast between tumor and stroma, both elasticity (Fig. 6c) and Euler angle (Fig. 6d) images present improved contrast that accentuates the boundaries of tumor regions with localized higher elasticity and lower Euler angle in the tumor compared to the surrounding stromal regions. This is consistent with the results shown in Fig. 4 and Fig. 5, respectively. In addition, unlike elasticity, Euler angle imaging provides a meaningful value at each pixel in the FOV, providing more complete visualization of the tissue. For example, a large portion of elasticity values in the stroma in the top left of Fig. 6c are negative and are, therefore, masked from the image. These regions correspond to regions of elevated Euler angle in Fig. 6d. Similarly, a localized Euler angle approaching 90º is observed in several dispersed regions containing ducts with DCIS, corresponding to negative elasticity that is also masked from the elastogram. To further exemplify the complementary contrast of Euler angle, two regions containing stroma and IDC were selected, and are highlighted by the magenta and green boxes in Figs. 6a–6d. These regions are magnified and presented in Figs. 6e–6h. Furthermore, in Figs. 6i–6k, we demonstrate the corresponding histogram plots of OCT, elasticity, and Euler angle within the same regions. Firstly, based on image texture, the two regions can be readily distinguished in the elastograms of elasticity (Fig. 6g) and Euler angle (Fig. 6h), however, it is challenging to distinguish between the two regions based on OCT intensity (Fig. 6f). This is further illustrated in the corresponding histograms, which quantitatively show the closely overlapped OCT intensity distribution (Fig. 6i) of the stromal (*μ*_OCT, S_ = 24.4 dB) and IDC (*μ*_OCT, IDC_ = 24.7 dB) regions, The histograms of the elasticity and Euler angle show much better delineation between the regions, with two distinctive peaks detected in the elasticity distribution (Fig. 6j) (*μ*_E, S_ = 42.0 kPa; *μ*_E, IDC_ = 130.0 kPa) and Euler angle distribution (Fig. 6k) (*μ*_α, S_ = 42.5º; *μ*_α, IDC_ = 22.2º), corresponding to two different tissue types.

**Fig. 6.**
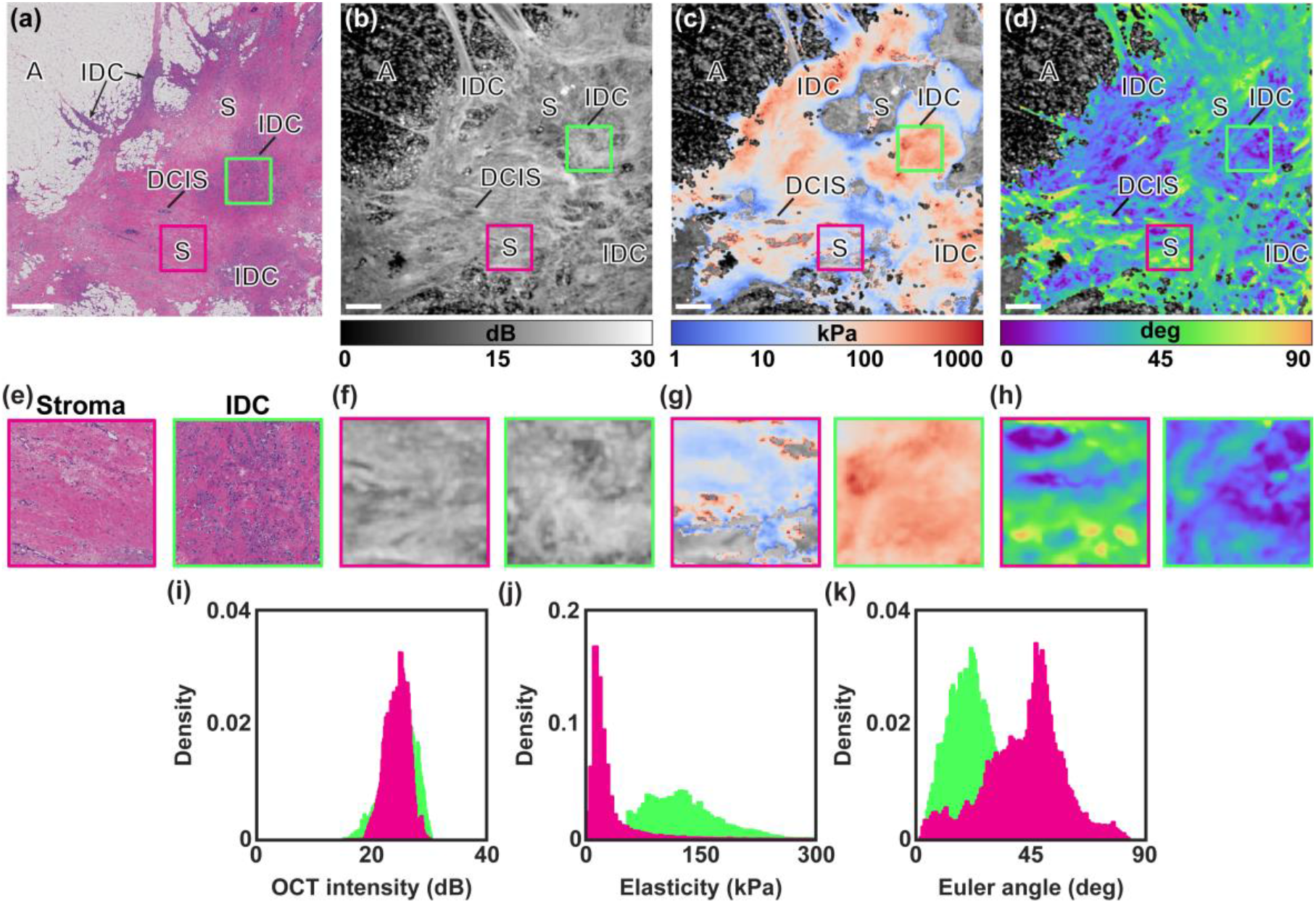
A mastectomy specimen containing IDC (Grade 2) and DCIS (intermediate grade). (a) The histology of the specimen. The co-registered *en face* (*xy*) images of (b) OCT, (c) elasticity, and (d) Euler angle, ∼140 µm below the tissue surface. The magenta and green boxes with an area of ∼1 mm × 1 mm in (a)–(d) indicate magnified regions in (e)–(h) corresponding to stroma and IDC, respectively. The magnified images represent (e) histology, (f) OCT, (g) elasticity, and (h) Euler angle, respectively. Histograms of the (i) OCT intensity, (j) selasticity, and (k) Euler angle corresponding to the magnified regions. A: adipose tissue; S: stroma; IDC: invasive ductal carcinoma, DCIS: ductal carcinoma *in situ*. Scale bars represent 1 mm.

In Figs. 7a–7c, we present boxplots of OCT intensity, elasticity, and Euler angle of regions of stroma (magenta) and breast tumor (invasive and non-invasive, green), respectively. Excluding one specimen that had poor co-registration with histology, nine mastectomy specimens were involved in the statistical analysis. As not every specimen contained both stroma and invasive tumor, two non-overlapping regions with an area of 0.5 mm^2^ were selected for at least one tissue type in each specimen, resulting in a total of 16 independent regions of stroma and 14 independent regions of tumor. The mean values of OCT intensity, elasticity, and Euler angle were then recorded in each region, corresponding to the scatter plots shown in Figs. 7a–7c, respectively. To evaluate whether OCT intensity, elasticity, and Euler angle are significantly different between stroma and tumor, a two-sample Kolmogorov-Smirnov test was implemented, which tests whether two populations come from the same distribution. The analysis shows no statistically significant difference (ns) between OCT intensity in stroma and tumor (*p* = 0.06 > 0.05), but significant differences between stroma and tumor, with a statistical significance of *p* = 4.1×10^−5^ < 0.001 (***) for both elasticity and Euler angle estimations.

**Fig. 7.**
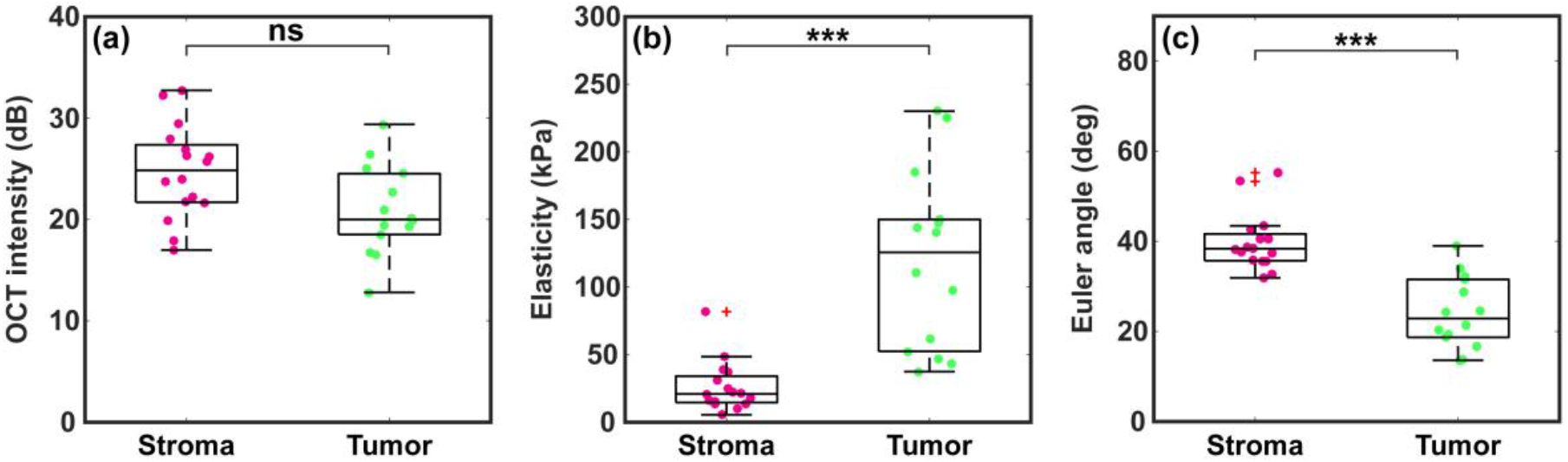
Boxplots illustrating the distribution of (a) OCT intensity, (b) elasticity and (c) Euler angle averaged across regions of stroma (magenta) and tumor (green) including IDC, ILC and DCIS, after co-registration between elastograms and histology images. A two-sample Kolmogorov-Smirnov test was used to assess whether stroma and tumor are from different populations, showing no statistically significant difference (ns) between the OCT intensity in stroma and tumor (*p* = 0.06 > 0.05), but statistical differences in both measurements of elasticity and Euler angle, corresponding to *p* = 4.1 × 10^−5^ < 0.001 (***). In each boxplot, the central mark indicates the median, the top and bottom edges of the box indicate the 25^th^ and 75^th^ percentile, respectively, and the top and bottom whiskers indicate the maximum and minimum values, respectively. The red addition sign indicates outliers.

## V. Discussion

Translation of OCE towards intraoperative tumor margin assessment requires both timely interpretation of elastograms and accurate identification of cancer. The assumption of uniaxial stress simplifies the mechanical model in OCE, facilitating rapid generation of elastograms. However, intrinsic tissue heterogeneity often invalidates this assumption, leading to erroneous elasticity estimation that must be masked from the image to avoid misinterpretation. In this study, we have shown that mapping the Euler angle of principal compression, calculated from the 3-D strain tensor, provides additional contrast to OCT and elasticity based on tissue heterogeneity. The enhanced contrast aids in distinguishing between invasive tumor and the surrounding stroma.

As shown in the elastograms (Figs. 4–6) and boxplots (Fig. 7) of breast tissue, regions of stroma tend to exhibit high Euler angle and low elasticity, whilst regions of invasive tumor tend to exhibit low Euler angle and high elasticity. The formation of unique patterns in these regions in Euler angle images is caused by the distinct deformation of the tumor and its microenvironment. Breast stroma is mainly composed of collagen-rich connective tissues [47]. The increased Euler angle of principal compression in stromal tissue, which highlights the increased deviation from the uniaxial compression, may also indicate the degree of mechanical anisotropy associated with the composition and organization of stroma, which can be remodeled during pathological development. For instance, the histology image in Fig. 4e reveals fibrotic stroma characterized by the proliferation of fibrous tissue with the removal of mammary acini and ducts [48]. The corresponding striations of high Euler angle in Fig. 4h highlight the distribution of stromal fibrosis. This is distinctive to the adjacent IDC tumor aggregates, which exhibit heterogeneous but relatively lower Euler angle, suggesting that the solid tumor mass is generally more palpable with mechanical heterogeneity. The corresponding elasticity measured in the region of tumor is higher than that of stroma, which is consistent with similar findings reported in previous studies [13], [14], [25]. In the future, integrating polarization-sensitive OCT into OCE would enable mapping of phase retardation and optic axis orientation within the tissue volume [49], [50]. This would provide validation of the measured Euler angle in stroma, which is more birefringent than invasive tumors [49], [50], and would also enable multiparametric quantitative characterization of breast cancer on the micro-scale for potentially improved diagnostic accuracy of tumor margin assessment.

This study was conducted using scans acquired with a phase-sensitive OCE system. This facilitates the ready use of a hybrid 3-D displacement estimation method, which combines phase-sensitive detection and a non-iterative complex cross-correlation method. The lateral and transverse components of displacement were estimated using the complex cross-correlation method with 5–10× lower spatial resolution than phase-sensitive detection [27], [38], due to the use of a sliding window (defined by a Gaussian kernel in this study). This limits the resolution of the strain tensor components, resulting in a mismatch in the resulting elastogram resolution between Euler angle and elasticity that is calculated from phase-derived axial strain and stress. For instance, the elastograms of ILC presented in Fig. 5 show that elasticity (Fig. 5g) preserves sharper tumor boundaries compared to the corresponding Euler angle image (Fig. 5h). This mismatch in spatial resolution could potentially be reduced by incorporating a more sensitive method to estimate non-axial displacement components, such as optical flow, which can provide displacement sensitivity up to 450× smaller than a single pixel [51] and has been combined with deep learning to improve the elastogram quality in ultrasound elastography [52]. Whilst phase-sensitive detection allows displacement to be measured only along the axis of the incident beam, it can be adapted for 3-D displacement estimation by illuminating the sample from different inclined angles [53]. One caveat is the increased system complexity with multiple independent spectral-domain OCT interferometers and light sources, which typically require careful and prolonged alignment.

In this proof-of-concept study to investigate a new OCE contrast mechanism to visualize breast cancer, we focused on breast tissue specimens excised from patients undergoing mastectomy. This simplified the identification of tumor, as the sections were close to the tumor mass and allowed elastograms in the *en face* plane to be directly compared with co-registered histology of the mastectomy specimens. A future study is needed to assess the diagnostic accuracy in tumor margin assessment by performing this technique on a larger number of breast-conserving surgery specimens, as has previously been performed for elasticity imaging in OCE [13].

## VI. Conclusion

In this study, we have presented novel mechanical contrast on the micro-scale based on tissue heterogeneity. We demonstrated the capability to differentiate invasive tumors from the surrounding stroma by mapping the Euler angle of principal compression into elastograms, which provides additional contrast to both OCT and elasticity. Our results show that the technique holds promise in the identification of malignant tumor during surgery, which could facilitate improved diagnostic accuracy of intraoperative tumor margin assessment.

